# BiasDB: A Comprehensive Database for Biased GPCR Ligands

**DOI:** 10.1101/742643

**Authors:** Christian Omieczynski, Trung Ngoc Nguyen, Dora Sribar, Lihua Deng, Dmitri Stepanov, David Schaller, Gerhard Wolber, Marcel Bermudez

## Abstract

G protein-coupled receptors transmit signals across membranes via interaction with intracellular binding partners. While there is an imprinted signaling profile for each receptor, biased ligands are able to shift intracellular pathways resulting in different recruitment profiles. We present the first comprehensive database of all literature-known biased ligands as a resource for medicinal chemistry and pharmacology. In addition to careful manual curation, we provide an analysis of the data. *BiasDB* is available at https://biasdb.drug-design.de/.

## Introduction

G protein-coupled receptors (GPCRs) are omnipresent in human tissues and are involved in virtually every physiological process rendering them highly important drug targets^1^. Although 35% of currently marketed drugs directly target GPCRs, many aspects of their complex signaling network remain elusive^2-4^. Since a single receptor can signal through several intracellular transducers and thereby triggers a set of different pathways (Figure 1), the clinical outcome of ligand-dependent receptor response strongly depends on the profile of activated pathways^5-8^. Whereas activation of one pathway might be therapeutically desired, other pathways could account for adverse drug reactions or contradict the clinical effect. Each GPCR shows a naturally imprinted signaling profile, which typically represents the effect of physiological ligands^9, 10^. Biased ligands (also referred to as functional selective ligands) can shift this signaling profile towards other pathways (Figure 1), providing a way to pharmacologically fine-tune GPCR signaling^5-8^.

**Figure 1:**
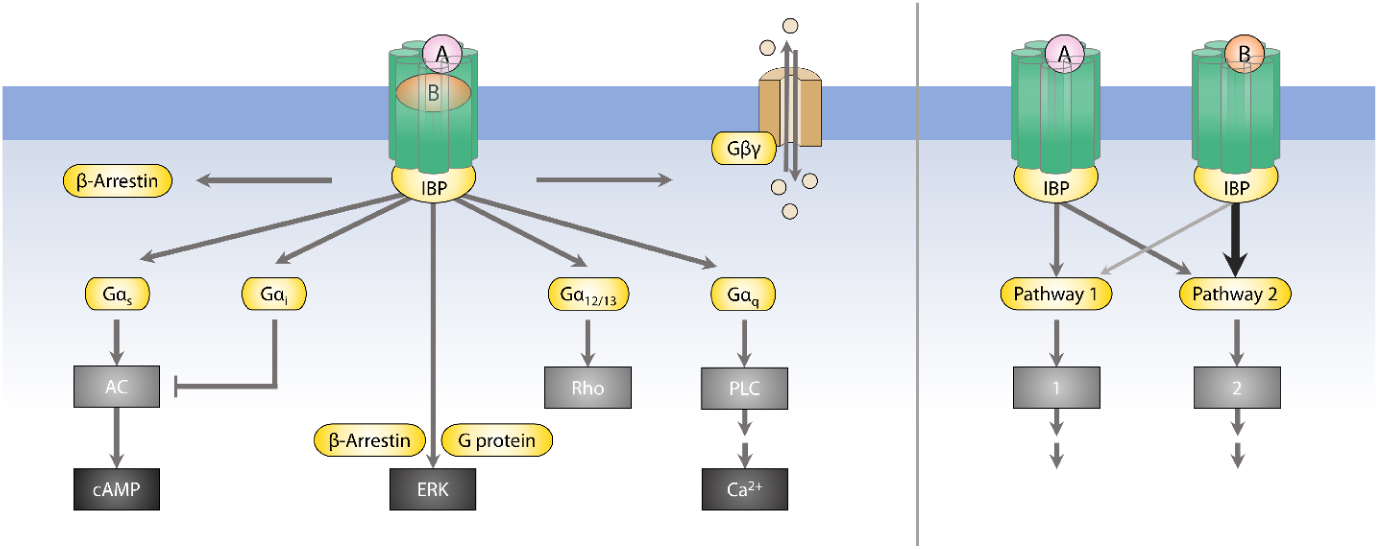
Simplified overview on GPCR signaling pathways and important effector proteins (left). Upon formation of the tertiary complex, which comprise a GPCR (green), ligand (A or B) and an intracellular binding partner (IBP), different signaling pathways can be activated through e.g. G proteins and β-arrestin (yellow), which can further trigger distinct effector proteins (grey). The concept of biased signaling in GPCRs involves a ligand-dependent shift of the activated downstream pathways (right). By taking Ligand A as a reference ligand, ligand B could be described as biased towards pathway 2.

In recent years, biased signaling has drawn more and more attention in the GPCR field, with many studies focusing on ligand design, assay development for bias determination and the resulting pharmacological outcome^5, 11, 12^. However, the structural prerequisites of biased ligands are poorly understood and only a few studies shed light on potential mechanisms for biased signaling^13-16^. Surprisingly, most biased ligands were discovered by either serendipity, extensive pharmacological testing or SAR studies based on known biased agonists^5^.

The importance of biased ligands as both tool compounds and drugs or drug candidates, demands a comprehensive overview on this class of ligands, but existing databases (e.g. ChEMBL or GPCRdb) lack information about signaling bias^17, 18^. Therefore, we systematically collected and manually curated data for the *BiasDB*, a database of known biased GPCR ligands as a resource for medicinal chemistry, chemical biology and pharmacology. Moreover, we provide a first analysis of the database content with regard to physicochemical properties and a comparison with clinically used GPCR ligands to identify potentially biased ligands.

## Results and Discussion

### Database Description

The *BiasDB* contains 618 cases of signaling bias representing 482 individual ligands for 61 receptors. We provide information about the chemical structure, target receptor, the type of bias, assay categories used for bias determination, the reference ligand, the literature source and standard molecular descriptors. Although we focused on small drug-like organic molecules, we also included peptide ligands with up to 13 residues. Within the *BiasDB* users can explore bias information by querying the above-mentioned criteria and moreover we provide a structure and similarity search. A snapshot from the website showing the organization of the user interface and a *BiasDB* scheme is given in the Supplementary Information.

An overview on the content of *BiasDB* is given in Figure 2. The ligand bias category was assessed in a hierarchical manner, in which we grouped bias types based on the preferred pathway, e.g. G protein bias contains several individual bias types such as G_i_/β-arrestin or G_s_/ERK. The vast majority of biased ligands are G protein-biased (56.8 %), followed by β-arrestin-biased ligands (24.6 %), ligands which show G protein selectivity (9.5 %) and ERK-bias (9.1 %). Interestingly, ligands with G_i_ over β-arrestin bias represent over one quarter of all bias cases (28.0 %). Not surprisingly, the number of reported biased ligands have dramatically increased over the last couple of years (Figure 2B) with aminergic GPCRs as the predominant target group. As expected, receptors which are widely used as model systems (e.g. D_2_, µ and β_2_ receptors) have a high number of reported biased agonists (Figure 2C). We would like to note that we have not included studies and ligands for which bias determination was not clear, since we don’t expect added value from these cases. This accounts for studies in which a reference ligand was missing, a known biased ligand was used as reference ligand, or the determined bias was not significant. We have not included quantitative bias data, since methods for ligand bias quantification are not comparable and a standardized approach is still missing in the field^7, 19^. We also excluded cases in which ligand bias was reported to be only time or tissue-dependent.

**Figure 2.**
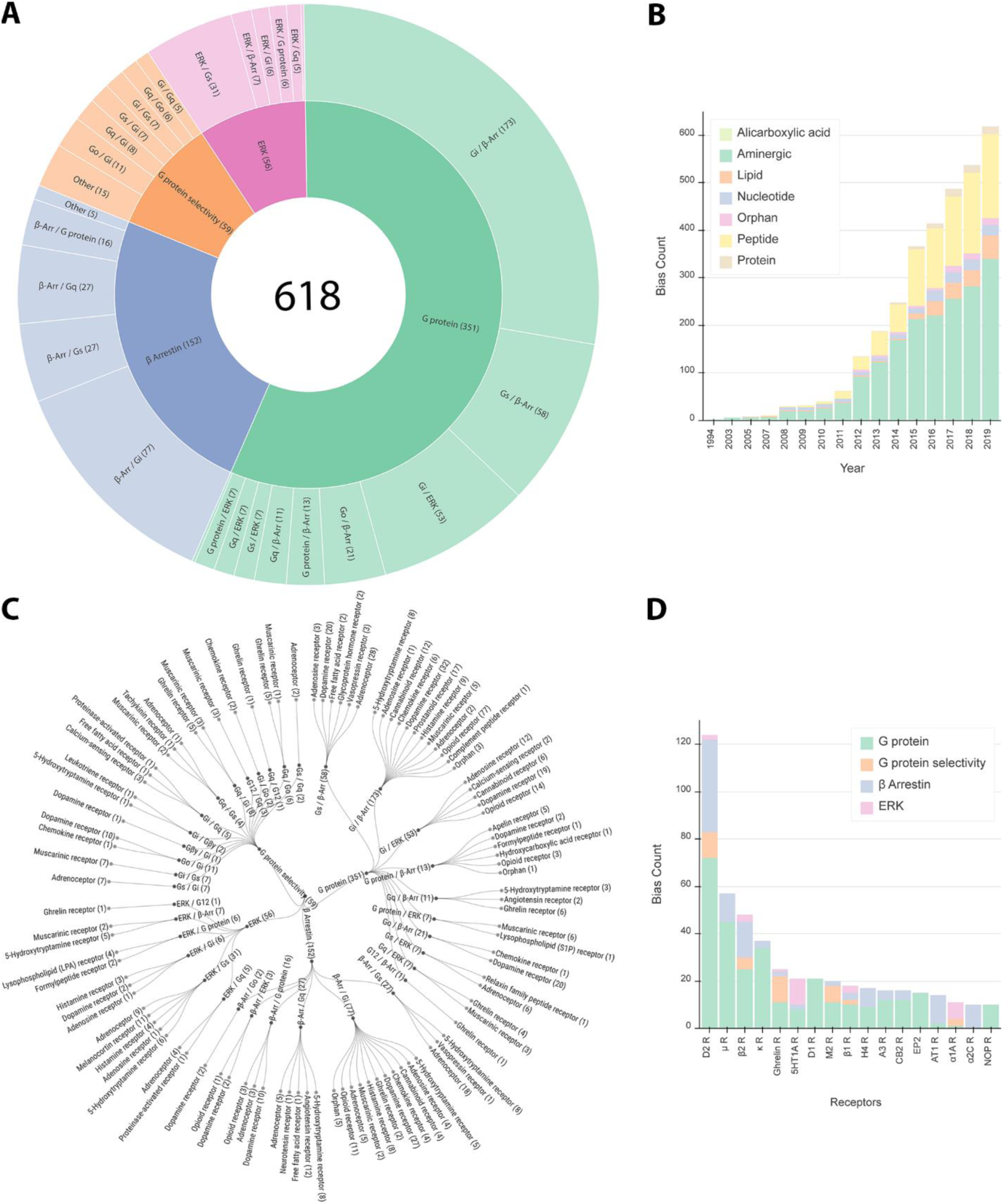
*BiasDB* data distribution in terms of bias type (A, groups with less than 5 entries were joined as ‘Other’), cumulated bias count per year (B), hierarchical overview on bias types (C) and the bias count of specific receptors (D, receptors with less than 10 cases are not displayed).

### Data Analysis

We calculated a set of six molecular descriptors (molecular weight, LogP, number of rings, number of hydrogen bond acceptors, hydrogen bond donors and topological polar surface area) for both the set of biased ligands in the *BiasDB* and their corresponding reference ligands to search for differences and trends in their molecular structure. The observed differences and trends might represent a good starting point for developing design strategies for biased ligands. The most prominent differences could be observed for molecular weight, LogP and the number of rings marking a tendency of biased ligands to be larger, more lipophilic and contain more rings compared to unbiased reference ligands (Figure 3A-C, Supplementary Information).

**Figure 3.**
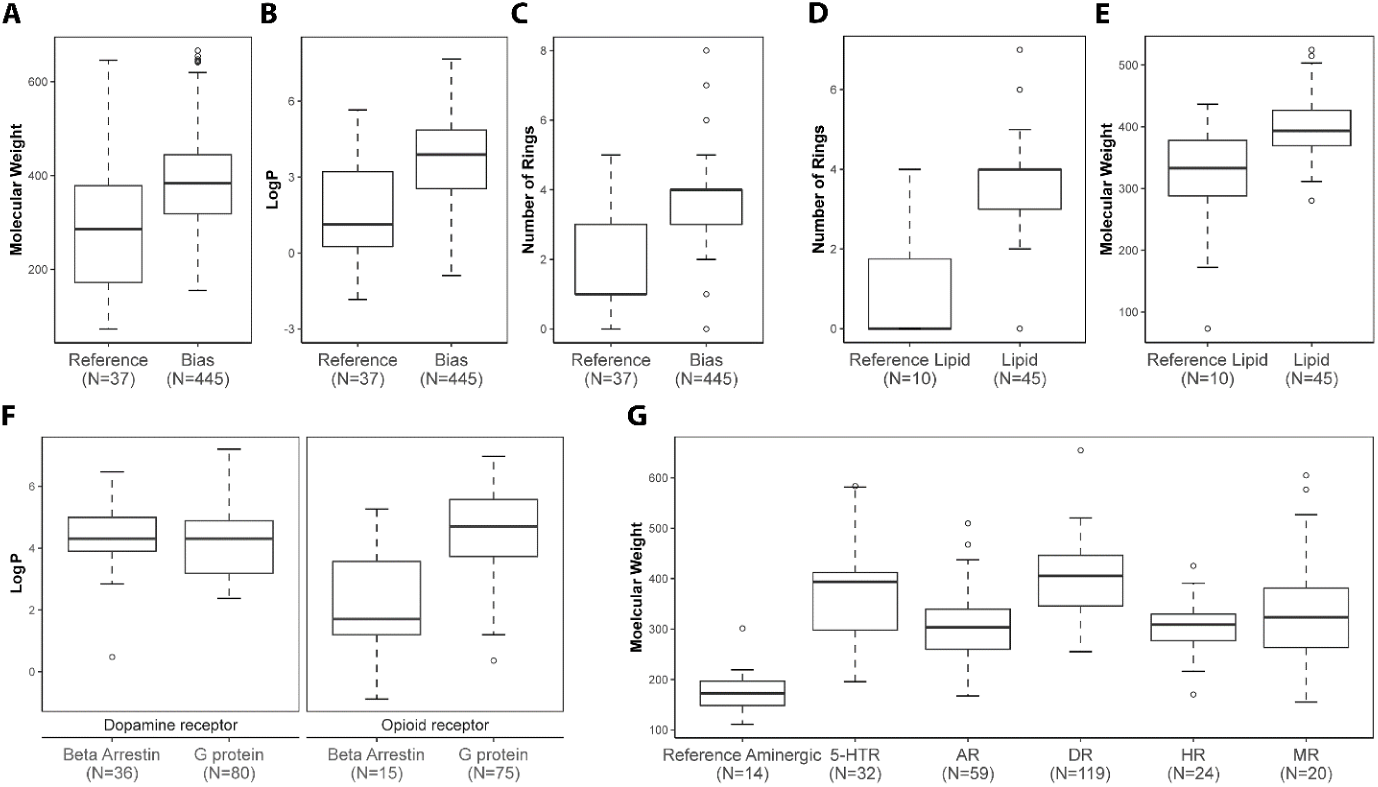
Chemical property analysis of *BiasDB* represented as box plots. Biased ligands show a general trend of having a higher molecular weight (A), being more lipophilic (B) and are composed of more rings (C) compared to reference ligands. Differences in property distribution for reference and biased ligands of for lipid receptors (D/E), or for different bias types for dopamine and opioid receptors (F) suggests a receptor family-specific pattern. The general trend shown in A-C, is even more pronounced for aminergic GPCRs.

These general trends have to be taken with caution, because they represent a mixture of ligands for different receptor types (e.g. aminergic, lipid or peptide binding receptors). We emphasize that different features might be helpful for different receptor types as exemplary illustrated for lipid receptors (Figure 3D-E). Whereas molecular weight seems to be less important, the number of rings might play an essential role for designing biased ligands for lipid receptors. However, since physiological ligands for lipid receptors are highly flexible due to their lipid nature, a common approach is to rigidify molecular structures to gain affinity. It is not clear whether the increased number of rings accounts for bias, or if this just reflects common trends in ligand design for lipid receptors. In another example, we looked for differences in molecular descriptors for biased ligands with a different bias category. We suggest that increased lipophilicity (LogP) of ligands might support G protein-bias versus β-arrestin-bias for opioid receptors but doesn’t play a crucial role for dopamine receptors (Figure 3F). Since aminergic GPCRs play an extraordinary role as drug targets, we expanded our analysis on different aminergic receptor families. We found a similar trend compared to the whole database regarding molecular weight, LogP and the number of rings. However, this trend was more pronounced for aminergic GPCRs (Figure 3F, Supplementary Information). This finding supports a recently reported concept for designing biased ligands by an extension of the molecular structure towards extracellular receptor regions^13^. We surmise that a large fraction of biased ligands for aminergic receptors are in line with this concept and facilitate their bias by conformational interference with the extracellular loop region. Interestingly, biased ligands for serotonin and dopamine receptors were found to be highly similar with respect to the applied descriptors and the observed trends were even more pronounced than for other aminergic GPCRs. The above-mentioned examples indicate that trends in physicochemical properties could guide synthesis-driven approaches, but receptor family and bias type must be taken into account.

### Potentially Biased Drugs

Since biased signaling is a relatively new phenomenon (Figure 2B) and nearly all currently marketed GPCR drugs were developed without taking signaling bias into account, it is tempting to hypothesize that a large fraction of these drugs show bias. However, little is known about potentially biased drugs in clinical use and only a few studies have addressed this issue^20, 21^. Therefore, we used a structural similarity approach to find marketed drugs which are likely to show biased signaling due to their structural similarity to known biased ligands. We combined a 2D similarity search based on Morgan fingerprints with a 3D shape-based approach. We identified molecule pairs of which one compound is a biased ligand and the other compound is a marketed drug with no reported bias (Figure 4). We found examples in which the molecular structure was enlarged by additional motifs (e.g. Levallorphan contains an allyl group instead of a methyl group in Levorphanol). In other examples ring structures contain more heteroatoms (e.g. pirbuterol contains a pyridine instead of a benzol ring like in salbutamol). Due to the high structural similarity, we surmise that there is a high probability that these drugs show biased signaling and point to the importance of a systematic pharmacological evaluation of marketed drugs with regard to biased signaling. Interestingly, we found many examples from different therapeutic areas and with different target GPCRs. The full list of molecule pairs can be found in the Supplementary Information. Assessing the bias properties of marketed drugs might help to mechanistically understand their clinical effect and their safety profile, in particular for pharmacological differences within a drug class.

**Figure 4.**
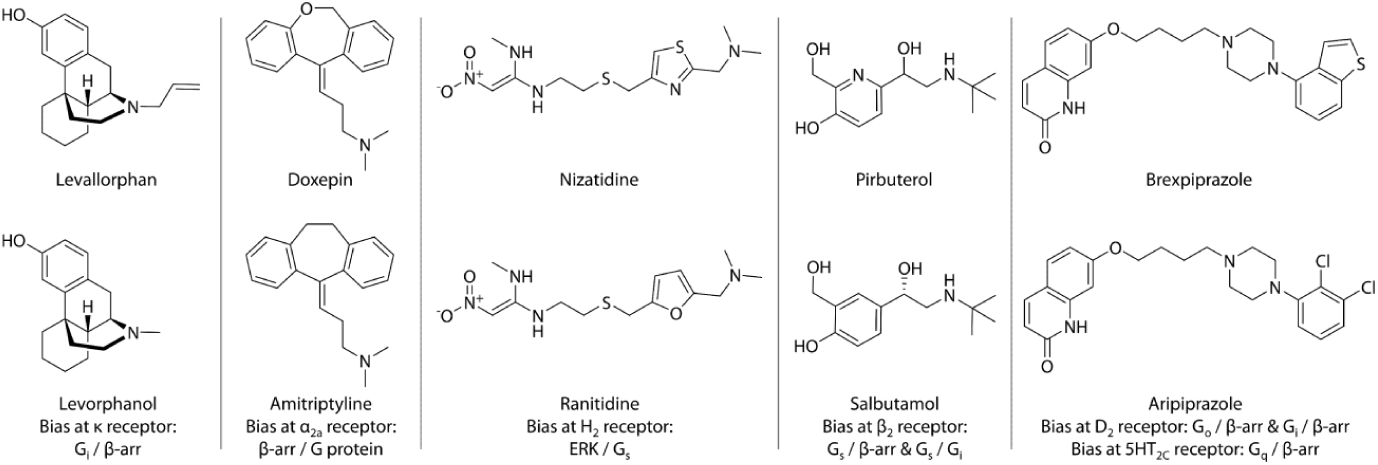
Potentially biased approved drugs from different drug classes (top row) found through similarity searches against *BiasDB* entries (below). The selected molecular pairs show only minor changes in their molecular structure and bind to the same respective receptors.

## Conclusion

Taken together, the *BiasDB* represents a novel resource for researchers in the GPCR field including medicinal chemists, pharmacologists and computational biologists, since it gathers information about biased ligands in a unique and comprehensive manner. Our first basic data analysis shows first insights into ligand properties linked to biased agonism, which could be helpful for rational ligand design. In particular, the recently suggested concept of *binding mode extension* is supported by our data analysis^13^. The molecule pairs identified by structural similarity emphasize that existing GPCR ligands are a likely source for biased agonists and require a systematic testing.

## Methods

### Data search and selection

The initial standardized search was executed with SciFinder by filtering GPCR related literature for either ‘ligand bias’, ‘biased agonism’, ‘biased signaling’ or ‘functional selectivity’. Furthermore, we complementary used PubMed and Google Scholar with the same search criteria and additionally searched for each receptor family separately covering the scientific literature till July 2019. Relevant bias information was extracted manually and carefully cross-checked.

### Database

*BiasDB* is based on a relational SQL database (MariaDB, https://mariadb.org/) (Supplementary Information), chemical information is processed using RDKit (https://rdkit.org/), structure searches are implemented using Chemaxons Marvin JS (https://chemaxon.com/products/marvin-js), 2D structure visualization uses kekule.js (http://partridgejiang.github.io/Kekule.js/). Visualization for the data was performed using D3 (https://d3js.org/) and google charts (https://developers.google.com/chart/). The web application is hosted as a Flask web application (https://palletsprojects.com/p/flask/) on a Linux server.

### Data analysis

Analysis of molecular descriptors was performed in R 3.5.1 (R: A language and environment for statistical computing. R Foundation for Statistical Computing, Vienna, Austria) using ggplot2. Molecular descriptors were generated using RDKit (http://www.rdkit.org). Molecules with molecular weight larger than 700 Da were excluded from small molecule analysis. Distributions for the analyzed molecular descriptors are represented as box plots, where the central line represents median of the data, and the lower and upper limits of the box are the first and third quartile, respectively. The whiskers extend up to 1.5 times the interquartile range from the lower and upper limits of the box to the furthest point within that distance. Data points beyond that distance are represented individually as points. Chemical structures of approved drugs were retrieved from DrugBank version 5.1.4^22^ totaling 2413 entries. Structures of biased ligands were retrieved from *BiasDB*. Both sets were filtered in MOE 2019.0101 (Molecular Operating Environment, Chemical Computing Group, Montreal, QC, Canada) for molecular weight below 700 Da to focus on small molecules resulting in 2232 approved drugs and 446 biased ligands. We excluded molecules containing no carbon atom and assigned protonation states at pH 7 using the molecule wash function in MOE 2019.0101. 2D similarity between the two ligand sets was calculated using Morgan fingerprints as implemented in RDKit nodes in KNIME 3.7.1 (http://www.knime.com). For 3D similarity assessment 25 conformations were generated per molecule using Omega 2.5.1.4^23^ with adjusted parameters (maxconfs=25, rms=0.8, ewindow=10, maxtime=30, enumNitrogen=true, flipper=true). ROCS 3.2.0.4^24^ was used to calculate 3D similarity between approved drugs and biased ligands with adjusted parameters (cutoff=1.0, mcquery enabled).

## Supporting information

Molecular pairs BiasDB/drugbank

## Author Contributions

The manuscript was written through contributions of all authors. All authors have given approval to the final version of the manuscript. ‡These authors contributed equally.

---

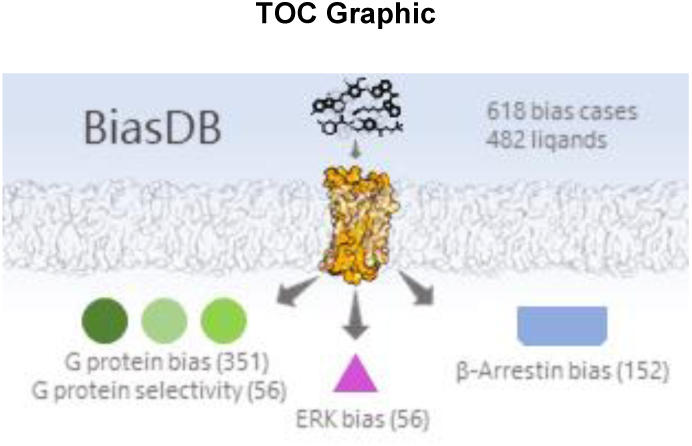

## Supporting Information for

**Figure S1.**
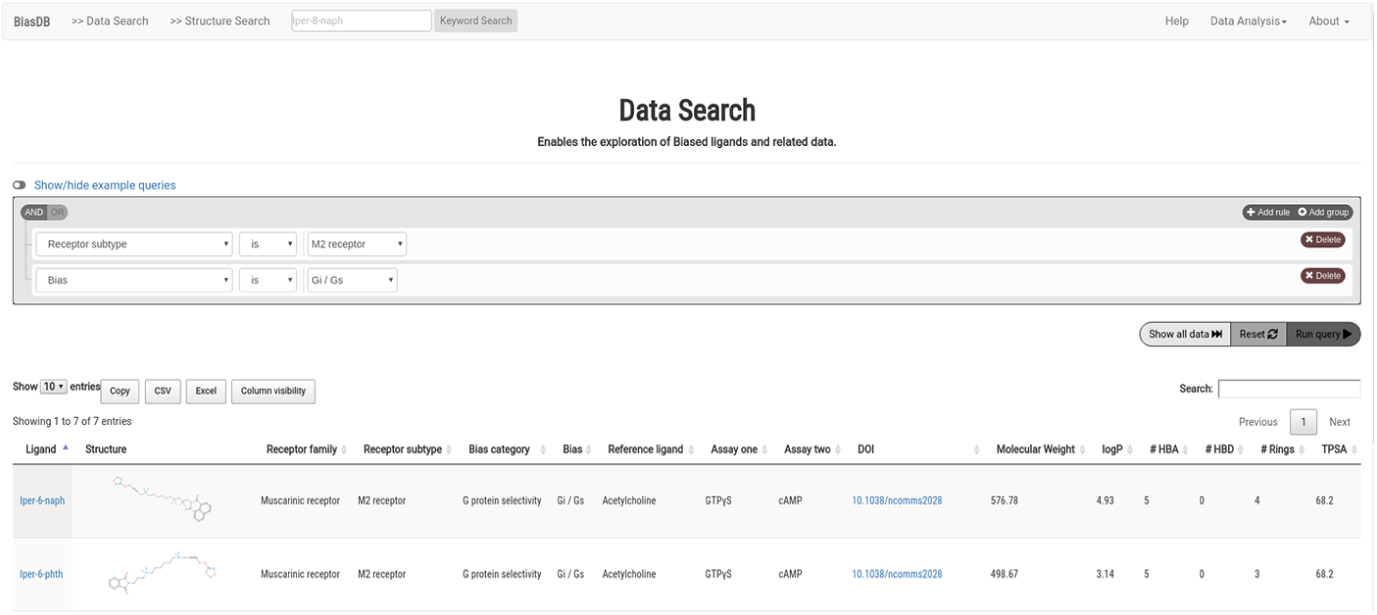
Screenshot of BiasDB web functionality. *By using queries in our “Data Search” the user can explore biased ligands and their data. We furthermore provide a full text search in the navigation bar and a “Structure Search”. Clicking on the results ligand name and structure will retrieve additional information.*

**Figure S2.**
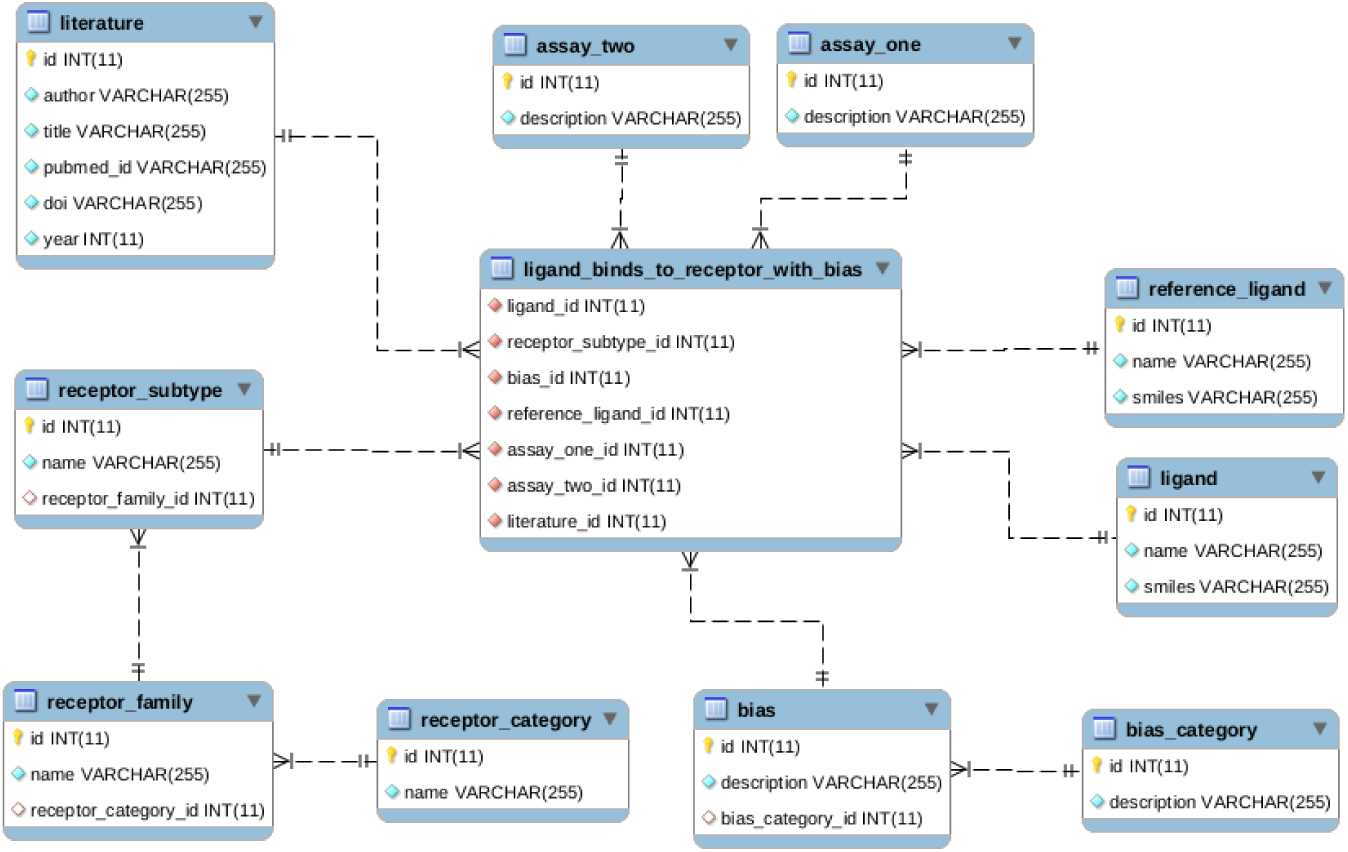
BiasDB scheme. All data tables are converging in the “ligand_binds_to_receptor_with_bias”-table, for presenting information about biased ligands. The “ligand”-table and “reference ligand”-table include structural information. The receptor information is hierarchically organized in three tables; “receptor_category”, “receptor_family” and “receptor_subtype” for retrieving data more easily. Information for bias is shown in two tables. The “bias_category” describes general bias such as G protein or β*-*Arrestin. The “bias”-table contains the actual bias, for example “Gs /Gi”. Additionally, information about used assays for bias detection and references is provided.

**Figure S3.**
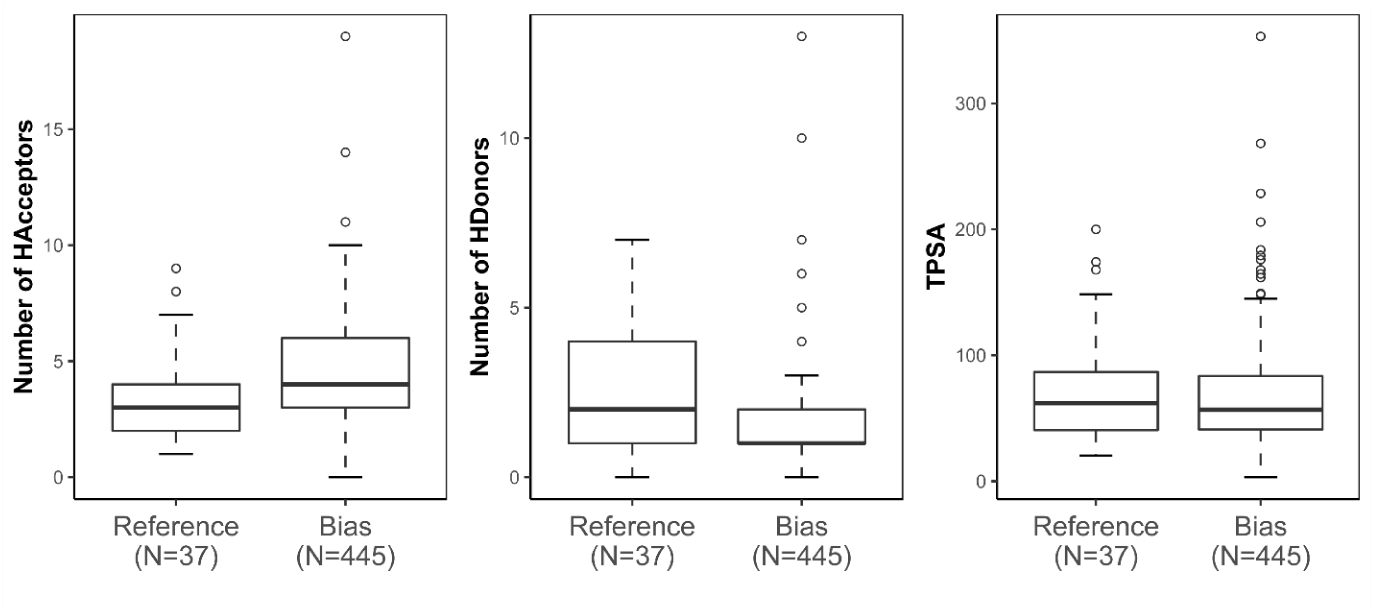
Comparison of the number of hydrogen bond acceptors, hydrogen bond donors and topological polar surface area (TPSA) between bias and reference ligand set.

**Figure S4.**
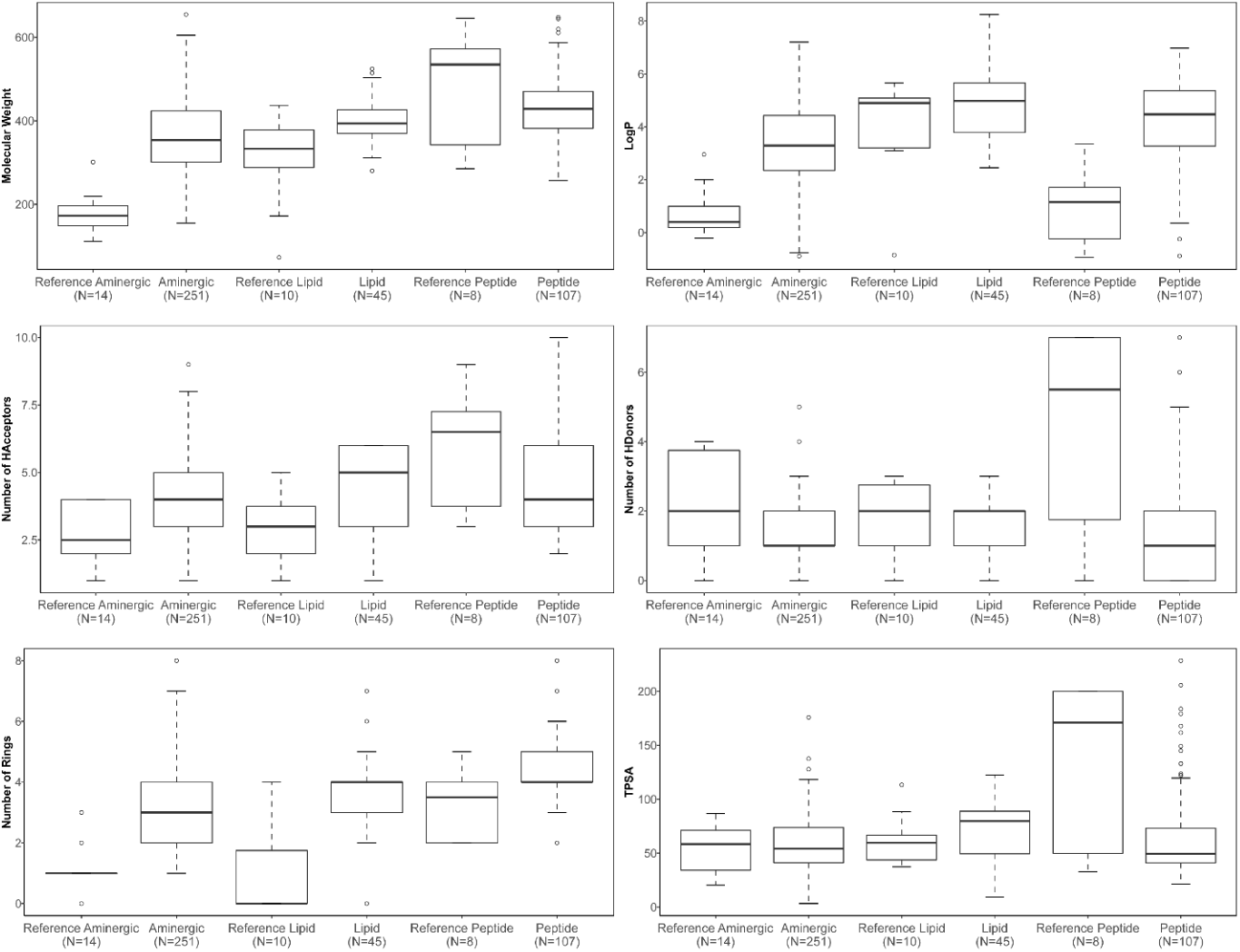
Comparison of 6 molecular descriptors (molecular weight, LogP, number of hydrogen bond acceptors, number of hydrogen bond donors, number of rings and topological polar surface area) between reference and biased ligands for the largest receptor categories (aminergic, lipid and peptide).

**Figure S5.**
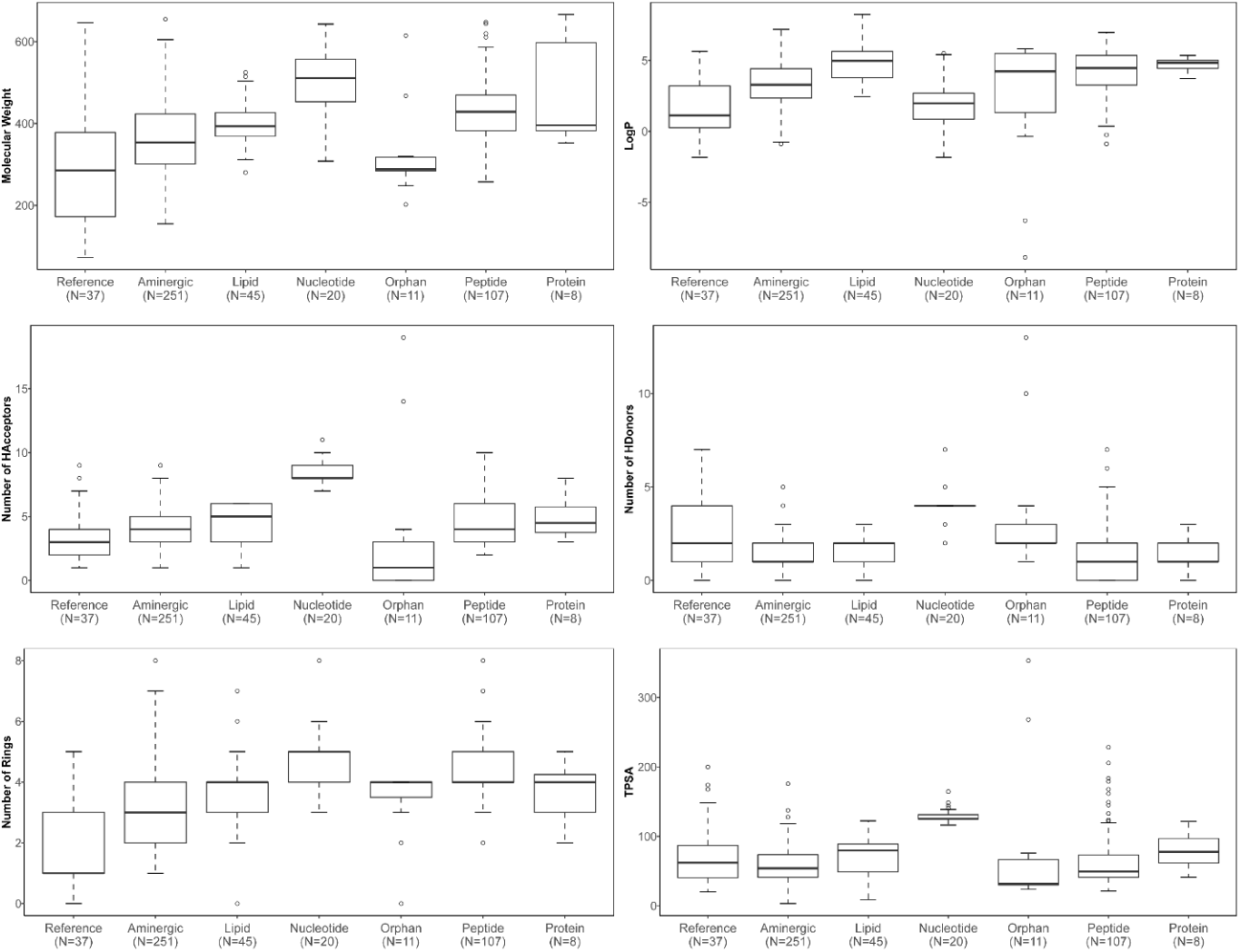
Comparison of six molecular descriptors (molecular weight, LogP, number of hydrogen bond acceptors, number of hydrogen bond donors, number of rings and topological polar surface area) between all reference and biased ligands for the specific receptor categories.

**Figure S6.**
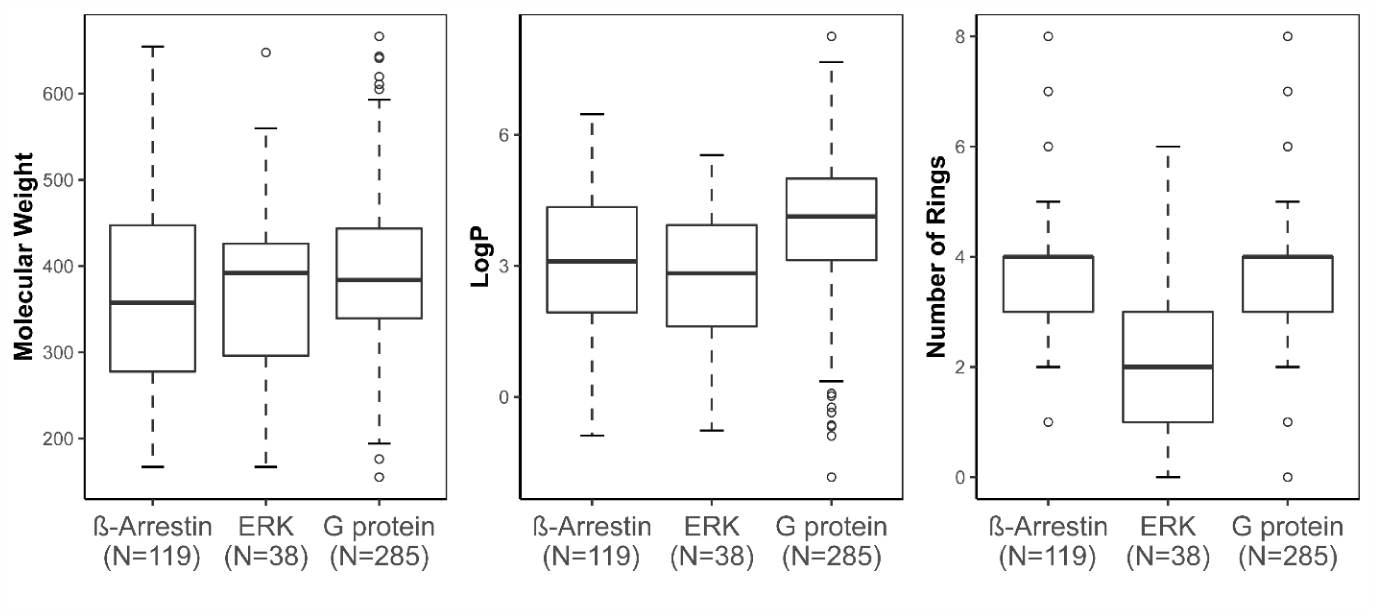
Comparison of molecular weight, LogP and number of rings for different bias types.

**Figure S7.**
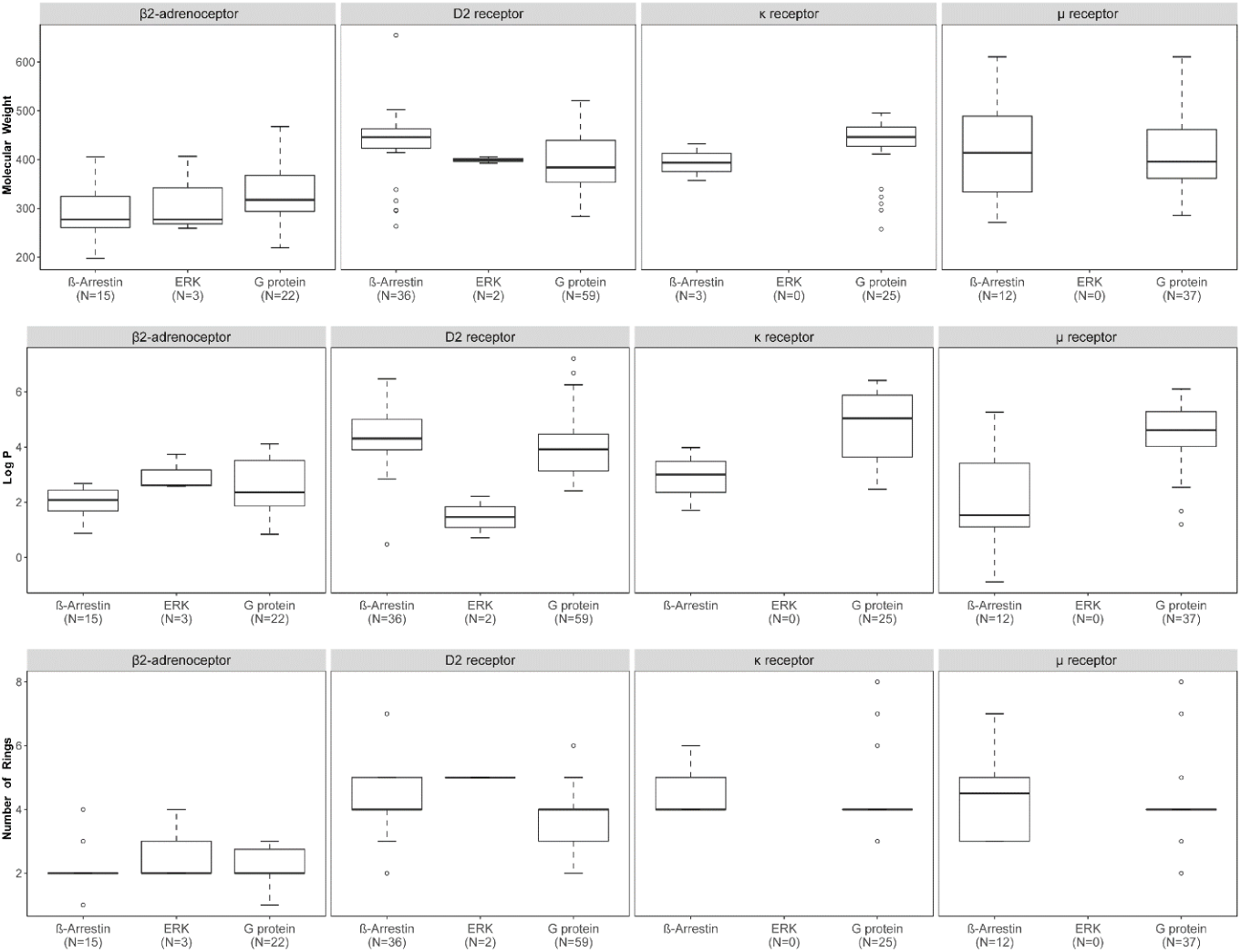
Comparison of molecular weight, LogP and number of rings for different bias types among different receptors subtypes.

**Figure S8.**
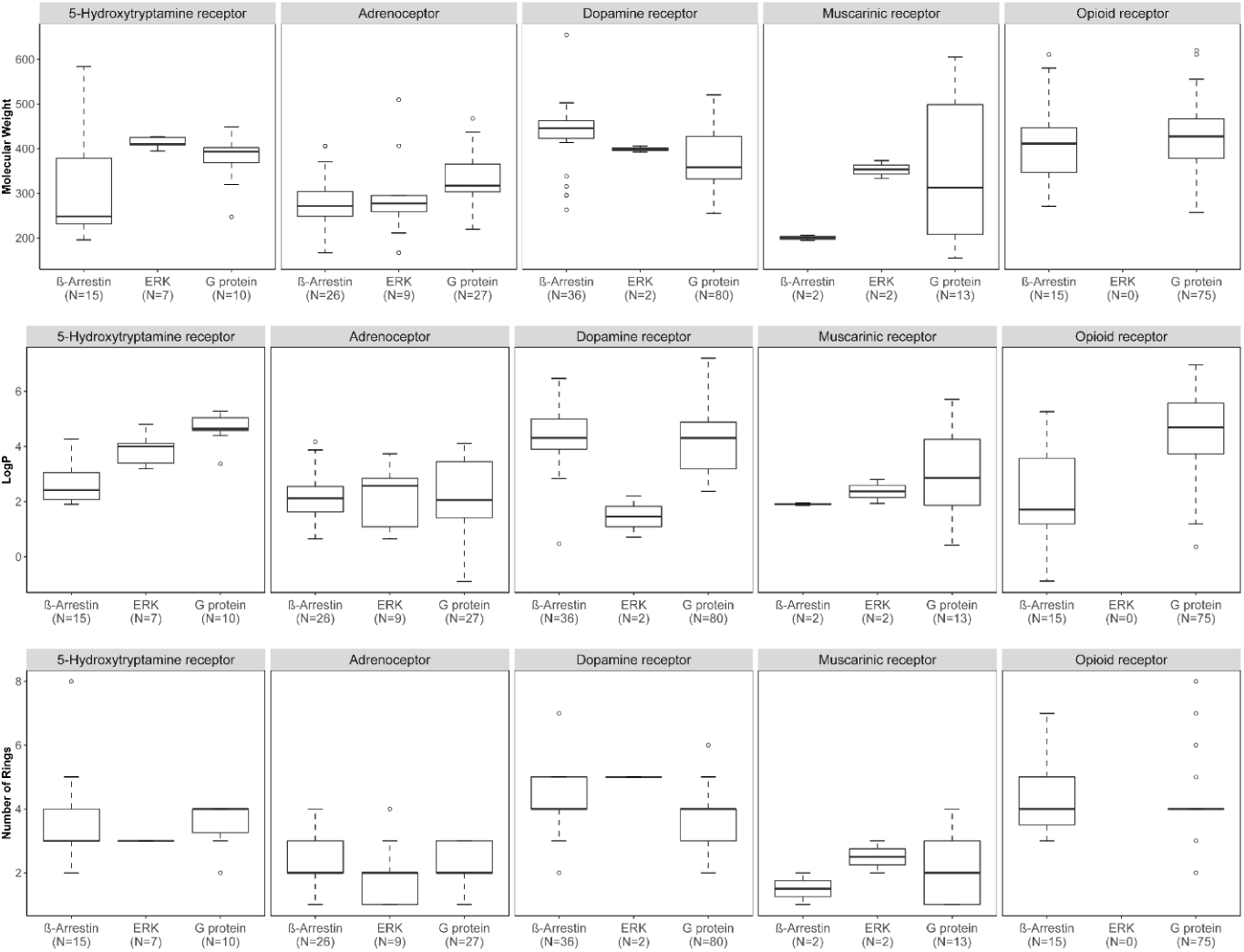
Comparison of molecular weight, LogP and number of rings for different bias types among different receptors families.

**Figure S9.**
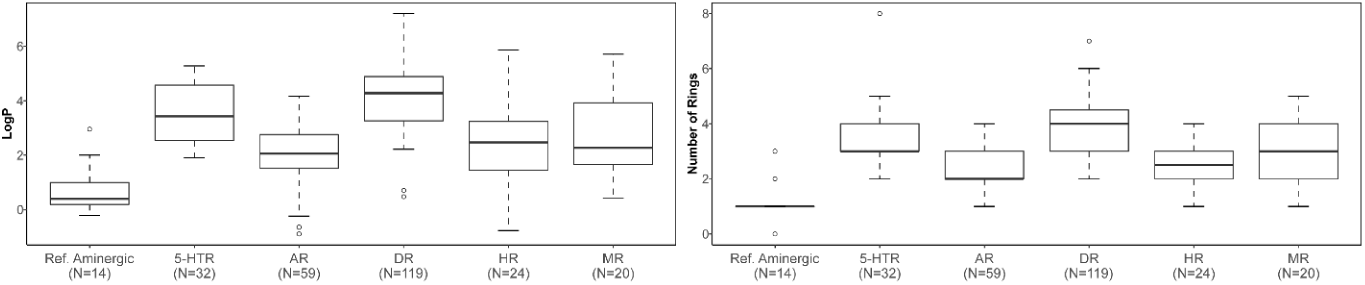
Comparison between LogP and number of rings between aminergic reference ligands and ligands for different aminergic receptor families. Abbreviations: 5HTR – 5-Hydroxytriptamine receptors, AR – Adrenoceptors, DR – Dopamine receptors, HR – Histamine receptors, MR – Muscarinic receptors

**Figure S10.**
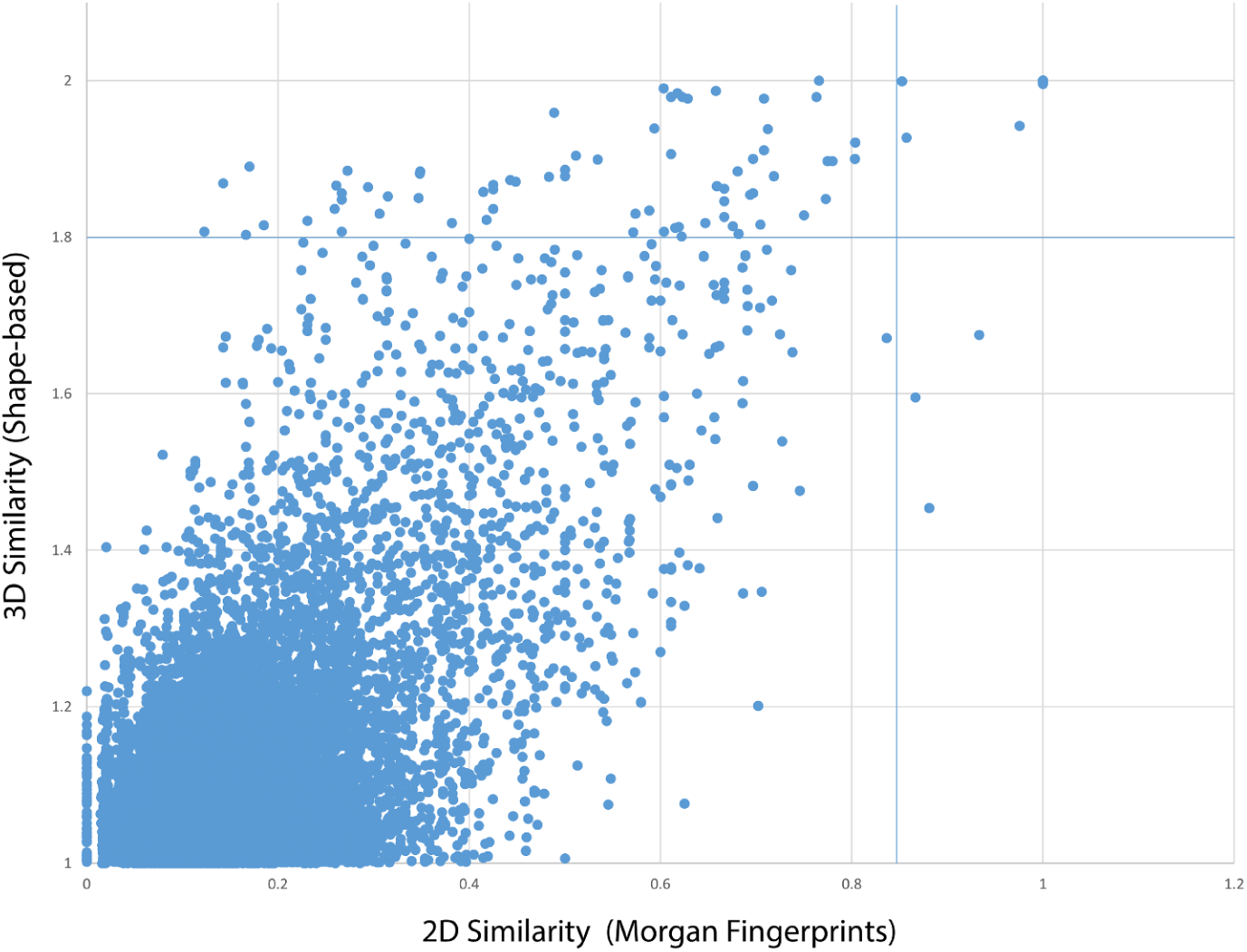
Distribution of molecular pairs based on 2D and 3D similarity. For 2D similarity we used Morgan fingerprints and for 3D similarity calculations we used a shape-based method with 25 conformations for each molecule. The molecular pairs in Figure 4 (BiasDB Paper) had thresholds of 1.8 3D similarity and/or 0.85 2D similarity (blue lines in Figure).

**Figure S11.**
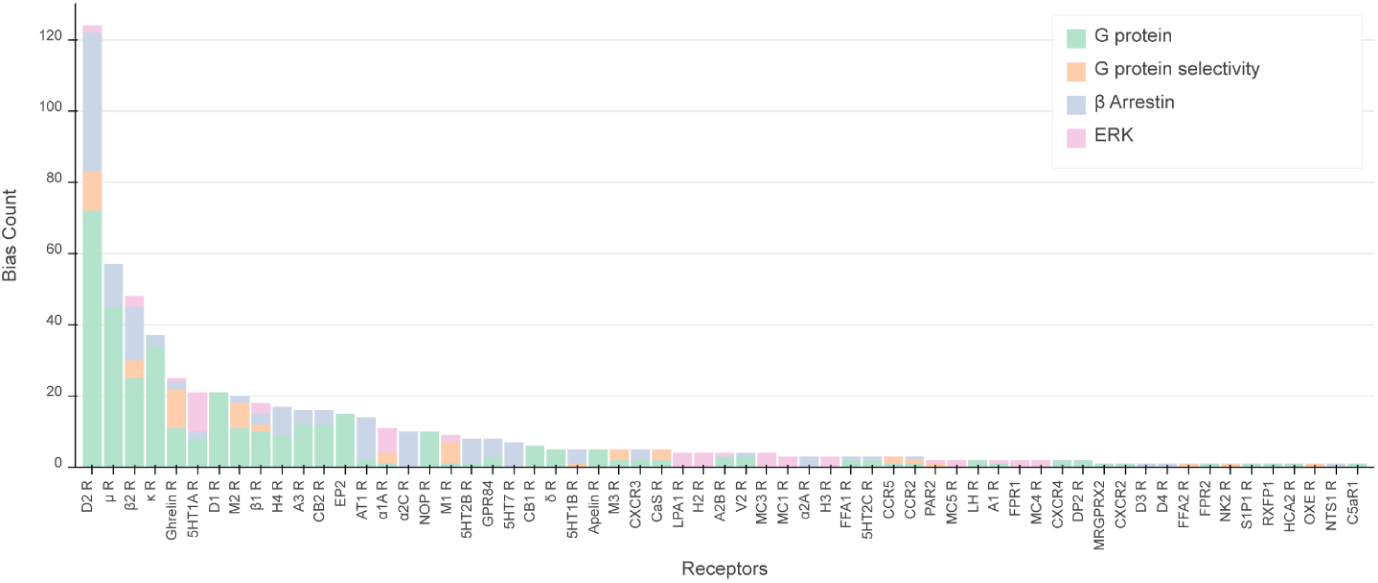
Full representation of biased ligands distribution over all receptors.

